# Psilocybin Reconfigures Cortical Dynamics Across Temporal, Spatial, and Hierarchical Scales

**DOI:** 10.64898/2026.11.22.768395

**Authors:** Donghui Song, Ze Wang, Henyi Rao

**Affiliations:** Department of Psychology, University of Science and Technology of China, Hefei, China; Department of Diagnostic Radiology and Nuclear Medicine, University of Maryland School of Medicine, Baltimore, MD, USA

**Keywords:** psilocybin, resting-state fMRI, brain entropy, cortical dynamics, surrogate analysis

## Abstract

Psychedelic fMRI studies report increased signal irregularity, but its spatial and temporal origins remain unclear. We quantified brain entropy (BEN) using sample entropy across space and time and tested BOLD surrogates in 57 paired PsiConnect participants. Both entropies increased after psilocybin, alongside reduced local synchrony, shorter intrinsic timescales, regional redistribution of low-frequency amplitude and contraction of a connectivity gradient. Surrogates retaining zero-lag voxel covariance reproduced the spatial entropy increase. Independent phase randomization preserving voxel power spectra reproduced the direction of temporal brain entropy change. Both measured increases and surrogate patterns recurred in an independent seven-person crossover cohort at 204 frames. Timescale maps corresponded at both scan lengths, while BEN maps corresponded after spatial correction at full duration. By linking brain entropy to spatial coordination and temporal persistence, this framework helps explain how psilocybin reorganizes cortical dynamics across scales and guides comparisons across pharmacologically altered brain states.

## Introduction

Acute psilocybin reshapes both the variability and coordination of spontaneous brain activity. Early fMRI work described a broader repertoire of transient connectivity states after psilocybin (Tagliazucchi et al., 2014). Later studies reported higher spontaneous MEG signal diversity across several psychedelics (Schartner et al., 2017) and higher fMRI time-series sample entropy after LSD (Lebedev et al., 2016). Precision fMRI localized acute psilocybin-associated desynchronization to distributed networks, including the default mode network (Siegel et al., 2024). PsiConnect further showed that changes in large-scale organization depend on the experiential context of dosing (Stoliker et al., 2026). These findings describe related aspects of the psychedelic state, from the succession of activity patterns to the coordination of signals across cortex.

The choice of entropy estimator determines which aspect of those changes becomes visible. A multimetric comparison found that the relation between acute psychedelic effects and fMRI entropy depends on the representation and estimator (McCulloch et al., 2026). Entropy over a sequence of connectivity states, for instance, concerns the repertoire or ordering of those states. Sample entropy concerns the probability that two similar motifs remain similar when extended (Richman & Moorman, 2000). Applied along the time axis of a voxel, it yields temporal brain entropy (BEN) and measures the recurrence of BOLD time-series motifs (Wang et al., 2014). Applied across adjacent voxels in one frame, the same recurrence rule yields spatial BEN (SampEn). Using both estimators on the same cleaned BOLD recordings makes it possible to ask which signal properties reproduce a change along each axis.

The temporal and spatial estimators suggest different statistical explanations. In a voxel, the power spectrum describes the distribution of variance across frequencies and encodes linear autocovariance. Shorter intrinsic timescales and larger successive-sample differences would make rapidly changing motifs more prevalent and motivate a spectral test of temporal BEN (Raut et al., 2020). Across voxels, correlations influence how often nearby BOLD intensities form similar within-frame patterns. Regional homogeneity characterizes local agreement across neighboring time courses (Zang et al., 2004), while summaries of cortical spatial modes and connectivity gradients characterize organization over longer distances. These complementary measures provide empirical context for testing serial dependence and cross-voxel covariance. Evidence that spatial and temporal autocorrelation can account for a range of functional network properties makes that distinction particularly relevant to fMRI (Shinn et al., 2023).

Surrogate signals make these explanations testable by preserving selected statistics before recomputing the same entropy measure. Applying the same Fourier phase shift to all voxels at each frequency retains their individual spectra and cross-spectra (Prichard & Theiler, 1994). Independent shifts preserve each voxel’s spectrum while disrupting the original cross-voxel phase relationships. A complementary rotation preserves the full zero-lag covariance matrix while reorganizing temporal order. A distance-dependent correlation model then tests how much of the spatial BEN change can be recovered from a common function of intervoxel distance. The comparisons quantify which statistics can reproduce the observed group mean changes under the tested constraints.

We characterized observed changes in entropy, local synchrony, temporal persistence and cortical organization in 57 paired PsiConnect participants (Novelli et al., 2026). We then tested which spatial and temporal signal properties reproduced the entropy changes under the surrogate models and assessed sensitivity to smoothing and global signal regression. Finally, we used an independent seven-person crossover cohort with methylphenidate as an active comparator to test whether the measured directions and surrogate patterns recurred (Siegel et al., 2024, Subramanian et al., 2025). Cross-cohort comparisons of regional effects at full and matched scan lengths asked whether a reproducible group mean response was accompanied by a reproducible cortical distribution.

## Results

### Study 1 Characterizing multiscale changes after psilocybin

Spatial BEN and temporal BEN both increased in PsiConnect (Fig. 1c,d). Spatial BEN increased in 49 of 57 participants, and the surface display localizes voxelwise FDR-significant BEN increases in association and visual cortex (Fig. 1a). ReHo showed the most extensive corrected decrease among the metrics in Fig. 1b, with decreases in 18,083 of 22,622 gray-matter voxels and in 18,207 after adjustment for changes in mean framewise displacement. Intrinsic timescales shortened by an average of 0.136 s, with corrected decreases in 4,320 voxels (Supplementary Fig. 1).

Higher mean squared successive differences (MSSD) in 9,649 voxels complemented the reduction in intrinsic timescale. The DFA exponent decreased in 7,678 voxels, and Lempel–Ziv complexity increased in 250, after false discovery rate correction. The amplitude of low-frequency fluctuations (ALFF) and its fractional form (fALFF) summarize different aspects of spectral amplitude. Their mean-normalized maps showed spatial redistribution. mALFF increased in 2,852 voxels and decreased in 4,884, while mfALFF increased in 2,485 and decreased in 2,585 (Supplementary Fig. 1). These amplitude measures contextualize the temporal response and motivate testing the full voxel power spectrum in Study 2.

**Figure 1.**
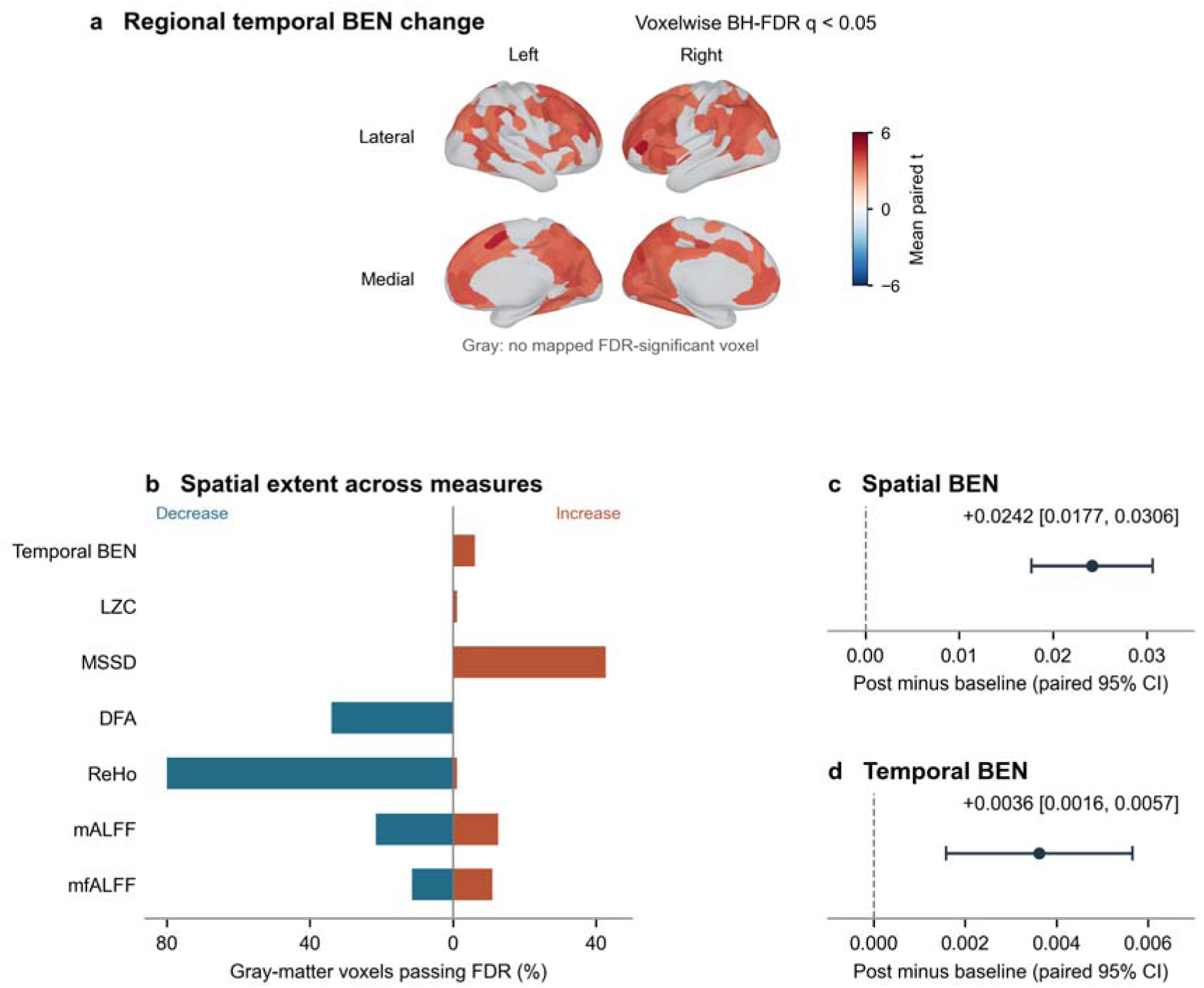
Observed changes in spatial and temporal BOLD dynamics in PsiConnect. (a) Inflated fsLR 32k cortex i the same 2 × 2 arrangement as Fig. 4, with lateral views above medial views and left and right hemispheres in the corresponding columns. Paired post-minus-baseline temporal BEN t values were first thresholded voxelwise within the analysis mask (BH-FDR q < 0.05). Colors show the mean signed t of passing voxels assigned to each Schaefer-400 parcel (Schaefer et al., 2018). Gray denotes regions with no mapped passing voxel. The display includes 1,34 of the 1,397 significant voxels. (b) Percentages of tested gray-matter voxels with corrected increases or decreases, including significant voxels outside the surface display, using each metric’s analysis mask as the denominator. (c,d) Mean paired changes and 95% confidence intervals for spatial BEN and temporal BEN in 57 participants, on separate axes.

The range and variance of the connectivity component most aligned with the sensory-to-transmodal axis decreased in PsiConnect (q = 0.0024 and 0.000845, respectively). The hierarchy-aligned absolute phase slope also decreased (q = 0.046). At the whole-cortex scale, voxelwise normalized global spatial complexity (NGSC) increased by 0.0192 (95% CI 0.0148 to 0.0235). NGSC and spatial BEN thus documented changes in two distinct spatial summaries, the distribution of variance across modes and the recurrence of local voxel patterns. Supplementary Fig. 2 shows the PsiConnect gradient contraction and the phase-slope change. Figure 4a shows the aligned Schaefer-400 gradient changes across cohorts.

We next asked whether the locations of the measured changes followed an independent cortical hierarchy or aligned with receptor reference maps. None of the eight effect maps showed a hierarchy association after false discovery rate correction. Six receptor–metric associations survived correction across all 224 molecular and cell-type comparisons. CB1 reference density aligned positively with MSSD and Lempel–Ziv changes and negatively with DFA and timescale changes. mGluR5 aligned positively with MSSD and negatively with DFA. The six associations also survived the fixed-mask spatial null. Spatial associations for the temporal profile persisted after adjusting for the independent hierarchy, while BEN associations with CB1 and mGluR5 did not survive that correction (Supplementary Figs. 3 and 4).

The two entropy increases thus occurred alongside changes in both spatial coordination and within-voxel dynamics. The surrogate comparisons separated these signal properties to test how much each could reproduce.

### Study 2 Testing which signal statistics reproduce the entropy changes

Reduced ReHo and higher NGSC implicated changes in spatial organization across scales. To test the role of cross-voxel structure in spatial BEN, we held individual power spectra fixed while varying whether the phase randomization preserved cross-spectra. We then tested how much of the spatial entropy response survived when the preserved spatial information was reduced to full zero-lag covariance or to distance-dependent correlations.

Shorter intrinsic timescales and higher MSSD pointed to changes in serial dependence. The DFA, Lempel–Ziv and low-frequency amplitude results extended that profile to scaling, sequence diversity and regional spectral amplitude. For temporal BEN, the key comparison was between independent phase randomization, which retains each voxel’s power spectrum, and zero-lag covariance surrogates, which retain contemporaneous spatial structure while disrupting the original time structure.

Spatial BEN increased under common phase randomization and full zero-lag covariance, while independent phase produced a small decrease (Fig. 2a). Across all 57 participants, full covariance recovered 99.6% of the common-phase group mean increase. Observed minus covariance was −0.00036 (95% CI −0.00149 to 0.00077), and common phase minus covariance was 0.00010 (95% CI −0.00002 to 0.00021). The same ordering held among the 49 participants outside the pilot, for whom covariance minus independent phase was 0.02443 (95% CI 0.01697 to 0.03188).

Both spectrum-preserving models reproduced the direction of the observed BEN increase, and the zero-lag covariance estimate was near zero (Fig. 2b). Independent phase minus covariance was 0.00532 (95% CI 0.00371 to 0.00693) across all 57 participants and 0.00484 (95% CI 0.00308 to 0.00660) among the 49 outside the pilot. The independent-phase point estimate exceeded the observed change, with observed minus independent phase at −0.00170 (95% CI −0.00342 to 0.00002, Fig. 2d).

The distance model recovered 97.4% of the full covariance model’s spatial BEN group mean increase across all 57 participants. The paired difference was 0.00063 (95% CI −0.00118 to 0.00244, Fig. 2a, c). The corresponding ratios were 97.3% in the 49 participants outside the pilot and 95.0% in the unsmoothed eight-person pilot. The paired intervals for both subsets spanned zero (Fig. 2c).

**Figure 2.**
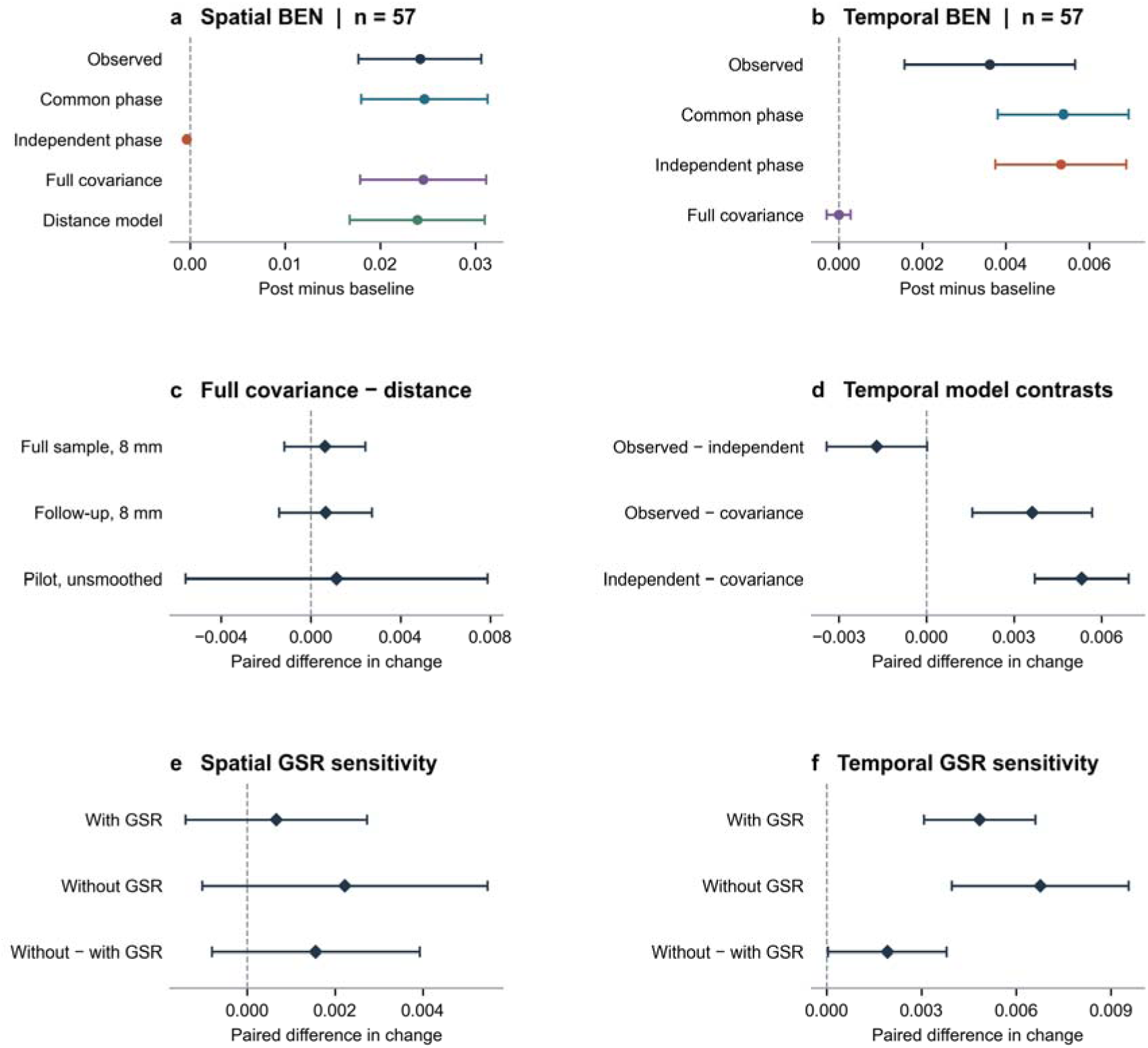
Signal structures that reproduce the entropy changes in PsiConnect. (a,b) Mean post-minus-baseline changes across 57 participants for observed signals and surrogate models. (c) Paired spatial BEN difference in change (full zero-lag covariance minus distance model) in the full sample (n = 57), the subset outside the pilot (n = 49) and the unsmoothed pilot (n = 8). (d) Selected temporal BEN contrasts in the full sample. (e,f) Sensitivity t global signal regression (GSR) in the same 49 participants, for full covariance minus distance (spatial BEN) an independent phase minus full covariance (temporal BEN). The bottom row in each sensitivity panel compares the contrast estimates within participants across pipelines. Points are group means and bars are t-based 95% confidence intervals. Each surrogate estimate averages 10 realizations per scan.

Omitting GSR preserved the spatial and temporal patterns in the 49 participants outside the pilot (Fig. 2e,f). The distance model reproduced 89.4% of the mean spatial BEN increase under full covariance, compared with 97.3% with GSR in the same participants. Full covariance minus distance was 0.00222 (95% CI −0.00103 to 0.00548) without GSR and 0.00066 (95% CI −0.00141 to 0.00273) with GSR. The within-participant shift after omitting GSR was 0.00156 (95% CI −0.00081 to 0.00393). For temporal BEN, independent phase minus zero-lag covariance was 0.00676 (95% CI 0.00396 to 0.00956) without GSR and 0.00484 (95% CI 0.00308 to 0.00660) with GSR. The paired shift after omitting GSR was 0.00192 (95% CI 0.00005 to 0.00379).

These contrasts identify a spatial covariance signature and a within-voxel spectral signature to test externally. The independent crossover data also allow us to distinguish recurrence of a group mean response from recurrence of its cortical distribution.

### Study 3 Testing external replication and active-comparator responses

#### Directional replication and surrogate responses

Spatial BEN increased and ReHo declined in all seven participants at both scan lengths, with NGSC increasing in six of seven at each length. In the common-support analysis at 204 frames, temporal BEN increased in all seven (Fig. 3b). The broader duration analysis showed higher Lempel–Ziv complexity and lower DFA in all seven at 204 frames, with higher MSSD in six. Intrinsic timescales shortened in five participants at full duration and seven at 204 frames. Supplementary Table 1 summarizes the metric-specific duration analysis.

In the 204-frame psilocybin-minus-baseline contrast, common phase and zero-lag covariance recovered the positive spatial BEN response, and the independent-phase point estimate was slightly negative (Fig. 3a). The full covariance model recovered 99.6% of the observed spatial group mean increase. Observed minus covariance was 0.00008 (95% CI −0.00147 to 0.00164), and covariance minus independent phase was positive in six of seven participants (group mean 0.02544, 95% CI −0.00279 to 0.05368).

For temporal BEN, the psilocybin-minus-baseline change was 0.00793 (95% CI 0.00237 to 0.01350) in the observed data and −0.00028 (95% CI −0.00096 to 0.00040) under zero-lag covariance (Figure 3b). The common- and independent-phase models both reproduced the positive direction. Their increases exceeded the covariance-model increase by 0.01146 (95% CI 0.00195 to 0.02096) and 0.01123 (95% CI 0.00166 to 0.02081), respectively.

The external distance model reproduced 85.6% of the full covariance model’s spatial BEN increase after psilocybin relative to baseline, below the 97.4% point estimate in PsiConnect. The external mean change was 0.02008 (95% CI −0.00299 to 0.04314) under the distance model and 0.02346 (95% CI 0.00124 to 0.04568) under full covariance. Their paired difference, full covariance minus distance, was 0.00338 (95% CI −0.00153 to 0.00829) (Figure 3a). For methylphenidate minus baseline, the distance-model change was 0.00867 (95% CI −0.00048 to 0.01783) and the model difference was 0.00046 (95% CI −0.00542 to 0.00635), with a group mean ratio of 94.9% (Figure 3c). For psilocybin minus methylphenidate, the corresponding estimates were 0.01140 (95% CI −0.00990 to 0.03271) and 0.00292 (95% CI −0.00324 to 0.00907), with a group mean ratio of 79.6% (Figure 3e). All three paired model-difference intervals spanned zero. The ratios describe group mean changes and are especially uncertain for the active-comparator contrasts.

Methylphenidate showed partly overlapping directions relative to baseline (Fig. 3c,d). For example, its independent-phase BEN increase exceeded the zero-lag covariance point estimate by 0.00675 (95% CI −0.00001 to 0.01351). In the direct drug comparison, the observed mean differences were 0.01496 for spatial BEN (95% CI −0.00667 to 0.03659) and 0.00170 for temporal BEN (95% CI −0.00537 to 0.00877). The observed and surrogate condition-difference intervals also spanned zero for both metrics (Fig. 3e,f).

**Figure 3.**
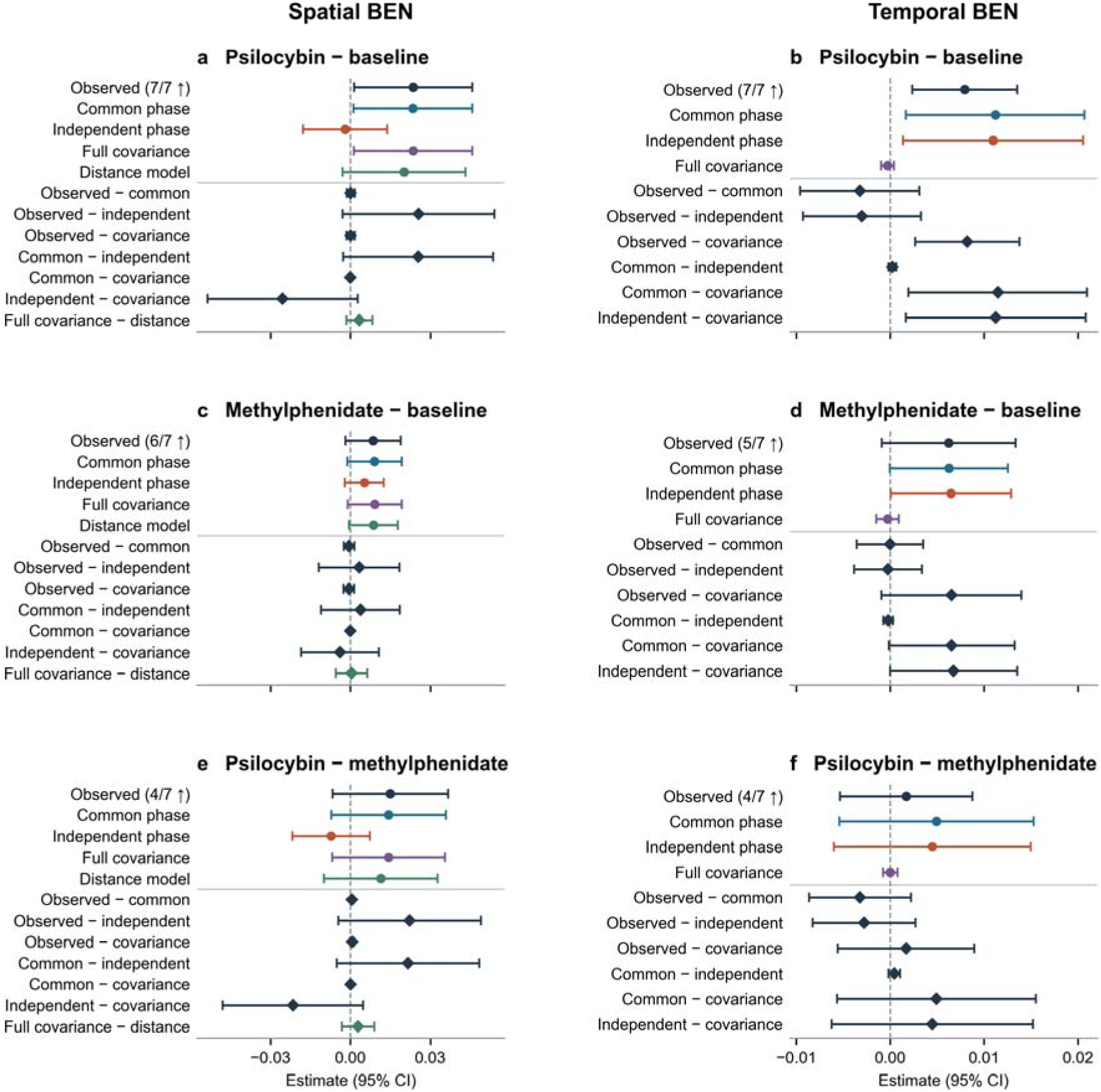
External replication of observed and surrogate entropy responses in ds006072 at 204 frames. (a,b) Psilocybin minus baseline. (c,d) Methylphenidate minus baseline. (e,f) Psilocybin minus methylphenidate. Circles show condition differences by signal type. In the observed row, the upward arrow denotes the fraction of participants with a positive difference. Diamonds below the thin horizontal divider show within-participant differences between the condition contrasts for two signal types. Points are means across seven participants and bars are t-based 95% confidence intervals. Each surrogate estimate averages 10 realizations per scan. Horizontal scales are shared within each metric.

#### Spatial correspondence of regional changes

The cortical surface maps show where temporal BEN and intrinsic timescale changed (Fig. 4a). Timescale shortening involved posterior cingulate and prefrontal territories, and its maps corresponded across cohorts at full duration and at 204 frames (Spearman ρ = 0.465 and 0.407, respectively). BEN showed regional correspondence at full duration (ρ = 0.379, spatial permutation q = 0.0073). At 204 frames its group mean still increased, while regional correspondence was weaker (ρ = 0.170, q = 0.182, Fig. 4b).

Regional amplitude changes also showed cross-cohort correspondence. The mALFF and mfALFF maps correlated across cohorts at both analysis lengths under the spatial null, complementing the mixed directions of the voxelwise amplitude changes. FC gradient maps were compared using a common reference. G3 corresponded at both lengths for psilocybin relative to baseline, and G1 and G2 corresponded at 204 frames (Fig. 4b). Methylphenidate relative to baseline also showed G3 correspondence. The external gradient comparison tests where connectivity coordinates changed, while the independent hierarchy analysis tests whether the effects follow an established cortical axis.

**Figure 4.**
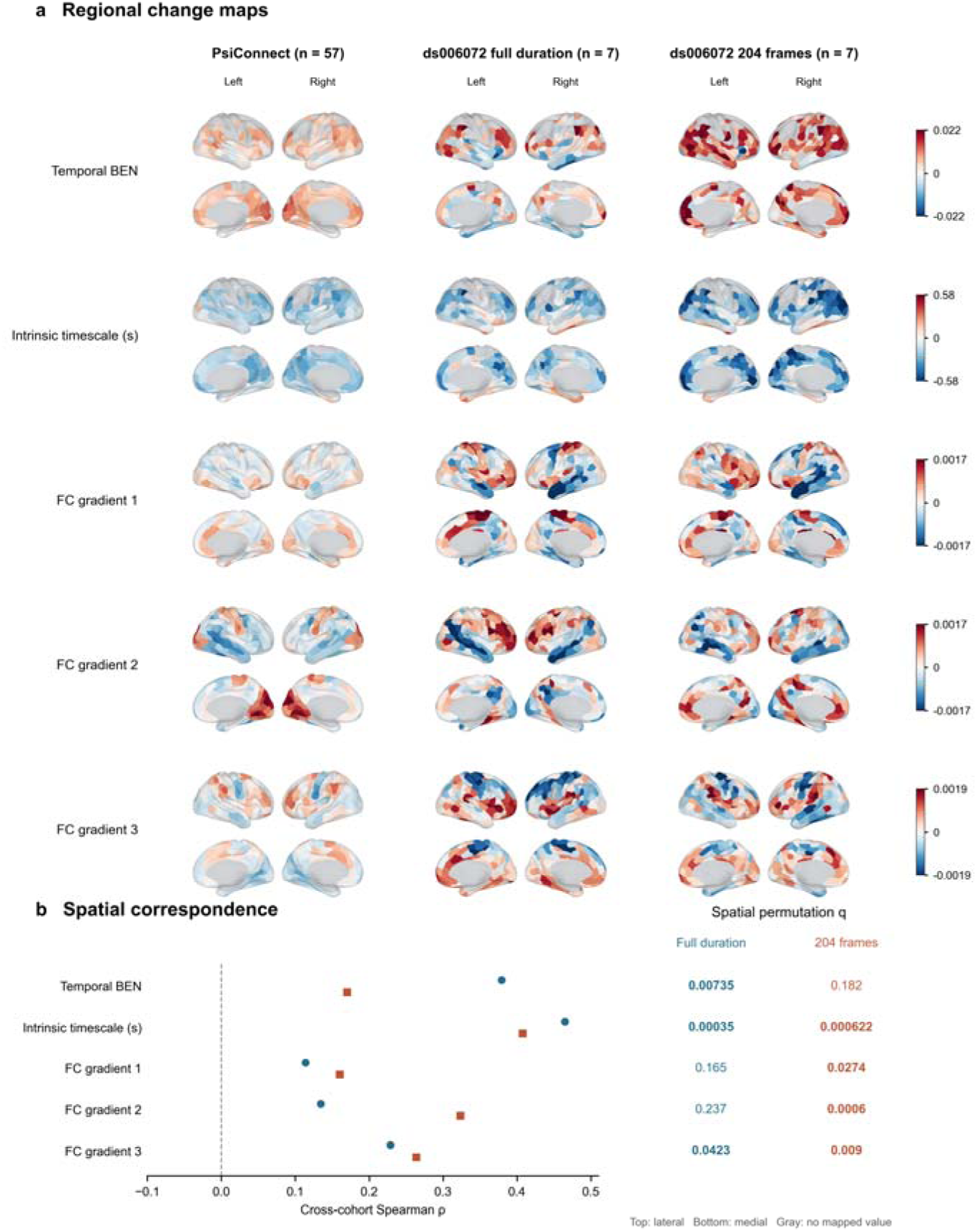
Cross-cohort correspondence of regional psilocybin-minus-baseline responses. (a) Schaefer-400 changes i temporal BEN, intrinsic timescale and three aligned functional connectivity gradients displayed on inflated fsLR 32 cortical surfaces. Each group is arranged in a 2 × 2 grid, with lateral views above medial views and left and right hemispheres in the corresponding columns. Red and blue indicate increases and decreases, respectively. Gra denotes unlabelled cortex or parcels outside common coverage. Maps are unthresholded. Each row uses a shared, zero-centered scale with limits at the pooled 98th percentile of absolute changes. (b) Spearman correlations between PsiConnect and ds006072 maps at full duration and 204 frames in ds006072. Bold q values indicate q < 0.05 after correction of the spatial permutation tests.

## Discussion

The paired PsiConnect scans showed that the same BOLD recordings can become more irregular along two different axes. Spatial BEN and temporal BEN both increased, while local synchrony decreased and intrinsic timescales shortened (Fig. 1). The surrogate comparisons resolve what these parallel increases capture. Spatial BEN retained its group mean increase when the surrogates retained contemporaneous correlations, and temporal BEN retained its positive direction when the surrogates retained individual voxel spectra (Fig. 2). The distinction puts earlier reports of psychedelic desynchronization and estimator-dependent entropy changes into a common measurement framework (Siegel et al., 2024, Stoliker et al., 2026, McCulloch et al., 2026).

Spatial BEN asks whether two within-frame patterns that match across three adjacent voxels still match when a fourth voxel is included. The probability of that extension depends on the joint distribution of nearby intensities. In a Gaussian field, the mean and covariance specify those joint distributions, so the probability of a spatial match can change even when the sequence of frames is reorganized. The phase comparison makes this concrete: common and independent phase randomization both preserve each voxel’s spectrum, and common phase also preserves the cross-spectra (Prichard & Theiler, 1994). The spatial increase persisted with common phase and was close to zero under independent phase. The full zero-lag covariance model recovered the common-phase increase while disrupting the original temporal ordering. Together, those comparisons identify within-frame covariance as a signal property sufficient to reproduce the observed group response. The decrease in ReHo and increase in NGSC show related changes in spatial coordination, from neighborhood concordance to the distribution of variance across cortical modes.

The distance model narrows that result. It retained voxel means and variances and replaced pair-specific correlations with a fitted function of intervoxel distance. It recovered 97.4% of the full covariance model’s spatial increase in PsiConnect and 85.6% in the external 204-frame contrast (Figs. 2 and 3). Spatial autocorrelation can account for features of resting-state connectivity that are often interpreted at the network level (Shinn et al., 2023). Here, a compact description of spatial correlation similarly captured much of the change in the recurrence of local patterns. The percentages compare group mean increases from two models. In the external cohort the paired full-minus-distance interval included zero and also allowed a larger contribution from pair-specific covariance. The unsmoothed pilot and the matched no-GSR comparison extend the spatial result across preprocessing choices, while the recovered fraction varies with the pipeline.

Temporal BEN extends a match forward by one observation within a voxel. Its match probability depends on the distribution of successive samples, which is shaped by serial dependence (Richman & Moorman, 2000, Xiong et al., 2017). At fixed variance, larger successive-sample differences are consistent with weaker dependence at lag one, while the shorter intrinsic timescales indicate reduced persistence over a broader interval (Fig. 1 and Supplementary Fig. 1). The power spectrum encodes circular autocovariance across lags. By holding spectra fixed while independently randomizing voxel phases, we recovered the positive BEN direction without requiring preservation of the original cross-voxel phase relationships (Fig. 2). The zero-lag covariance surrogate, which disrupts the original time dependence, yielded a near-zero mean change. Omitting GSR increased the independent-phase versus covariance contrast in the same 49 participants, preserving its direction. These results connect the BEN increase to the organization of power over frequencies. ALFF and fALFF locate regional changes in low-frequency amplitude and relative amplitude, and DFA and Lempel–Ziv complexity describe finite-scale scaling and sequence diversity. Together they contextualize the spectral result without assigning the BEN increase to a single frequency band.

The independent cohort tests how far these findings generalize. Relative to baseline, psilocybin increased spatial BEN and reduced ReHo in all seven participants at both scan lengths. In the common-support 204-frame analysis, temporal BEN also increased in all seven, and both spectrum-preserving models reproduced its positive group direction (Fig. 3). Intrinsic timescale shortening was present in five participants at full duration and seven using 204 frames. These observations reproduce the joint profile of higher irregularity and lower persistence despite differences in acquisition and sampling. Methylphenidate showed partially overlapping baseline-relative responses. The intervals for the direct psilocybin–methylphenidate contrasts spanned zero, so the external evidence supports recurrence of the psilocybin–baseline responses, with uncertainty about their differentiation from the active comparator.

A reproducible mean response can coexist with variation in where it is strongest. Using spatial rotations to account for autocorrelation in the Schaefer-400 maps (Alexander-Bloch et al., 2018), we found that the intrinsic timescale effect had positive spatial correspondence across cohorts at both scan lengths (Fig. 4). The maps showed recurring reductions in regions including posterior cingulate and prefrontal cortex. BEN corresponded at full duration, whereas at 204 frames the group mean increase remained evident and the regional correlation was smaller and did not survive correction. The window comparison therefore provides stronger anatomical support for reduced persistence than for the precise location of higher BEN. Spatial correspondence of MSSD, DFA, Lempel–Ziv complexity, mALFF and mfALFF at both durations adds evidence that the temporal response extends beyond any single estimator (Supplementary Results).

Connectivity organization supplies a further spatial scale. The sensory-to-transmodal component contracted in PsiConnect, and aligned FC gradient 3 showed regional correspondence at both external scan lengths, with gradients 1 and 2 also corresponding at 204 frames (Fig. 4). Component correspondence describes where connectivity coordinates changed. A separate test of the metric maps against an independent cortical hierarchy (Margulies et al., 2016) yielded no corrected associations. PET reference maps placed selected MSSD, DFA, Lempel–Ziv and timescale effects in regions with characteristic CB1 or mGluR5 distributions (Hansen et al., 2022). Some associations persisted after hierarchy adjustment, while neither receptor association with BEN passed correction in that analysis. The reference maps characterize the regional context of those effects using population-average receptor distributions.

The model contrasts concern the statistical organization of the BOLD signal. Psychedelic effects on hemodynamic responses and neurovascular coupling provide one reason to separate that inference from the physiology that generates the measured signal (Padawer-Curry et al., 2025).

The contribution is an empirical distinction within the recorded data: nearby voxel covariance can reproduce much of the spatial entropy response, individual spectra can reproduce the direction of temporal BEN, and cortical maps reveal which aspects of that temporal response recur regionally. Considering spatial and temporal sample entropy together thus links acute changes in irregularity to measurable properties of BOLD organization and to their generalization across cohorts.

## Methods

### Dataset and participants

Data were drawn from the publicly available PsiConnect dataset (OpenNeuro ds006110) (Novelli et al., 2026), an open-label multimodal study of psychedelic-naive adults scanned at baseline and after oral psilocybin administration (19 mg). Fifty-seven participants had paired baseline (ses-01) and post-psilocybin (ses-02) resting-state scans that passed the specified motion criterion: no more than 20% of volumes with framewise displacement greater than 0.5 mm in either session. This secondary analysis is a within-participant pre/post comparison without a placebo condition. The original PsiConnect study was approved by the Monash University Human Research Ethics Committee and registered as ACTRN12621001375842. Participants provided written informed consent.

### MRI data acquisition

PsiConnect (ds006110) acquired BOLD images at Monash Biomedical Imaging on a 3 T Siemens MAGNETOM Skyra. The T2*-weighted multiband, multi-echo echo-planar sequence used a repetition time (TR) of 910 ms, echo times of 12.60, 29.23, 45.86 and 62.49 ms, a multiband factor of 4, a field of view of 206 mm, right–left phase encoding and 3.2-mm isotropic voxels. The nominal eight-minute eyes-closed resting run contained 505 volumes. It was acquired first at each visit, with post-dose scanning starting approximately 80 min after administration (Stoliker et al., 2026, Novelli et al., 2026).

The external precision-mapping cohort (ds006072) was scanned at Washington University Medical Center on a 3 T Siemens Prisma. Resting-state BOLD used multi-echo echo-planar imaging with 2-mm isotropic voxels and 72 axial slices. TR was 1,761 ms, and echo times were 14.20, 38.93, 63.66, 88.39 and 113.12 ms. Multiband acceleration was 6, in-plane acceleration was 2, and the flip angle was 68°. Spin-echo images with opposing phase-encoding directions were acquired for distortion correction (Siegel et al., 2024, Subramanian et al., 2025). Run selection and duration matching for the present reanalysis are described below.

### Image preprocessing and denoising

We used fMRIPrep 22.0.2 (Esteban et al., 2019) preprocessed BOLD images in MNI152NLin2009cAsym space and the corresponding confound files. Confound regression included the Friston-24 expansion of six rigid-body motion estimates and the original, derivative, squared, and squared-derivative terms for global, white-matter, and cerebrospinal-fluid signals, together with an intercept and linear trend. Volumes with framewise displacement greater than 0.5 mm were interpolated after regression. Unless otherwise stated, signals were band-pass filtered at 0.01-0.08 Hz and standardized voxelwise. The first four cleaned volumes were discarded, leaving at most 501 analyzed time points per run. Analyses were restricted to a common gray-matter mask defined by tissue probability at least 0.30 and coverage in at least 80% of participants. Spatial analyses used 8-mm FWHM smoothing within the fMRIPrep brain mask; ReHo was calculated before smoothing and its final map was then smoothed by 8 mm. Although PsiConnect used multi-echo acquisition, this analysis used the fMRIPrep preprocessed BOLD series rather than a tedana-denoised multi-echo derivative.

### Dynamic measures

Temporal BEN used sample entropy with embedding dimension m = 3 and a tolerance of 0.6 times the voxel’s temporal standard deviation, excluding self-matches (Richman & Moorman, 2000, Wang et al., 2014). Regional homogeneity was Kendall’s W across a voxel and its 26 neighbors (Zang et al., 2004). Intrinsic timescale was the unbiased BOLD autocorrelation integral to its first zero crossing (Raut et al., 2020). Supplementary Methods detail additional measures. Supplementary Methods specify LZC, DFA, MSSD, mALFF, mfALFF, NGSC, connectivity gradients, hierarchy-aligned phase measures and molecular spatial inference.

### Global spatial measures

Spatial BEN was calculated within each BOLD frame from contiguous four-voxel templates along each of the three voxel axes. Match counts for three- and four-voxel prefixes used a maximum absolute difference threshold of 0.6 times the within-frame spatial standard deviation. Self-matches were excluded. We sampled 131,072 template pairs per axis, reused the same draws across conditions within each participant, and averaged the negative log ratio of four-to three-voxel matches over frames and axes. Spatial BEN used nuisance-cleaned, 8-mm-smoothed gray-matter BOLD in the primary analysis.

### Statistical inference

Paired comparisons used the participant as the unit of inference. Voxelwise p values were controlled within each gray-matter mask and metric with Benjamini–Hochberg false discovery rate correction. The surrogate analyses report t-based 95% confidence intervals for the paired differences in change. Additional inference procedures are described in Supplementary Methods.

### Independent cohort and matched duration analysis

We analyzed one baseline, one acute psilocybin and one acute methylphenidate resting run for each of seven participants from the independent randomized crossover dataset ds006072 (Siegel et al., 2024, Subramanian et al., 2025). We selected BOLDREST1 run 1 and used the first baseline except for P5, whose ses-3 baseline matched the phase-encoding direction of both acute scans. Open-label repeat visits were not included. Multi-echo BOLD was processed with fMRIPrep 25.2.5, including opposite phase-encoding distortion correction, and resampled onto the PsiConnect analysis grid. We used the same nuisance, smoothing and metric definitions with the native repetition time of 1.761 s. Within each participant and metric, condition means were computed over the intersection of valid gray-matter voxels across the three runs.

Full-length analyses used all available samples after the initial four-volume discard. The duration sensitivity analysis recomputed each metric using the first 204 retained frames (approximately six minutes), matching the shortest run. Nuisance regression and filtering were performed before window selection using the original metric-specific processing order; this analysis matches metric input length rather than fitting preprocessing anew on the short window. BEN used m = 3 and r = 0.6 times the temporal standard deviation of the analyzed series. We report paired difference signs for both drug-to-baseline contrasts and the direct psilocybin-minus-methylphenidate contrast.

### Covariance and Fourier surrogate analysis

We evaluated three surrogate models on cleaned, 8-mm-smoothed BOLD data from 57 paired PsiConnect participants after discarding the first four frames. Both Fourier models retained each voxel’s power spectrum, including its DC and Nyquist coefficients. Common-phase randomization used the same phase shift for all voxels at each frequency and retained the full cross-spectra (Prichard & Theiler, 1994). Independent-phase randomization used an independent shift for each voxel. For the zero-lag covariance model, we projected the centered signal into the orthonormal time subspace perpendicular to the constant vector, applied a random orthogonal rotation sampled by QR decomposition of a Gaussian matrix, and projected back before restoring each voxel’s mean. This generates new frames and preserves the sample voxel covariance matrix exactly, while redistributing power across frequencies and disrupting the original lagged dependencies. Each model used 10 realizations per scan.

Within each participant, the observed data and all surrogates shared the same gray-matter voxel support and spatial template draws. The pilot used the valid-voxel intersection across smoothed and unsmoothed scans. The 49 additional participants used the valid-voxel intersection across their two smoothed scans. Both entropy estimators used m = 3 and a tolerance multiplier of 0.6, applied to the temporal standard deviation for BEN and the within-frame spatial standard deviation for spatial BEN. We averaged each model’s realizations within scan and estimated paired post-minus-baseline changes and contrasts between those changes, using participant-level t-based 95% confidence intervals. We summarized the 49 participants outside the pilot separately and then pooled their estimates with the eight pilot participants.

In PsiConnect, we also modeled the spatial correlations as a function of intervoxel distance. For each scan, we fit a nonnegative mixture of six isotropic Gaussian correlation kernels and a zero-distance term to sampled correlations out to six voxel steps, giving greater weight to distances up to three steps. Kernel widths were fixed at 0.6, 1.0, 1.5, 2.3, 3.5 and 5.0 voxel steps. We generated independent Gaussian spatial fields by Fourier filtering on a padded lattice and rescaled each voxel to its observed mean and standard deviation. Each scan contributed 10 realizations evaluated on the same mask and spatial templates as the full covariance model. We report the mean within-participant difference between the two post-minus-baseline changes and its t-based 95% confidence interval. The reproduced fraction is the ratio of the group mean changes. The primary comparison used 8-mm data from the 49 participants outside the pilot and from all 57. We assessed the effect of omitting spatial smoothing in the original eight participants.

In the 49 PsiConnect participants outside the pilot, we repeated the focal spatial and temporal contrasts without global signal regression (GSR). We removed the global signal and its derivative and squared regressors from the nuisance model. All other regressors, motion interpolation, filtering, frame selection, voxel support, 8-mm smoothing, entropy parameters and template draws were retained. Each scan contributed 10 realizations per surrogate model. We computed post-minus-baseline changes for each participant, then compared full covariance minus distance for spatial BEN and independent phase minus zero-lag covariance for temporal BEN. The change in each model contrast between preprocessing choices was estimated within participant using t-based 95% confidence intervals.

We applied the same three surrogate constraints to the 21 preprocessed scans in ds006072. For each scan, we retained 204 consecutive frames after discarding the first four and used the same 8-mm smoothing, denoising, entropy parameters and spatial templates as in the external phase analysis. All three visits and four signal types for each participant shared valid gray-matter voxel support. Each surrogate estimate averaged 10 realizations per scan. We calculated psilocybin minus baseline, methylphenidate minus baseline, and psilocybin minus methylphenidate within participant for each signal type, then calculated pairwise differences between signal types. Confidence intervals were t-based and used the seven participant-level contrasts. We also applied the distance model to spatial BEN in all 21 external scans, using the same 204-frame windows, 8-mm smoothing, voxel support, template draws and 10 realizations per scan. For each condition contrast, we compared the paired change under full zero-lag covariance with the paired change under the distance model.

## Data availability

The original imaging datasets are publicly available from OpenNeuro under accessions ds006110 (PsiConnect, https://openneuro.org/datasets/ds006110) and ds006072 (Psilocybin Precision Functional Mapping, https://openneuro.org/datasets/ds006072).

## References

Alexander-Bloch AF, Shou H, Liu S, et al. On testing for spatial correspondence between maps of human brain structure and function. NeuroImage. 2018;178:540–551. doi:10.1016/j.neuroimage.2018.05.070.

Esteban O, Markiewicz CJ, Blair RW, et al. fMRIPrep: a robust preprocessing pipeline for functional MRI. Nature Methods. 2019;16:111–116. doi:10.1038/s41592-018-0235-4.

Hansen JY, Shafiei G, Markello RD, et al. Mapping neurotransmitter systems to the structural and functional organization of the human neocortex. Nature Neuroscience. 2022;25:1569–1581. doi:10.1038/s41593-022-01186-3.

Lebedev AV, Kaelen M, Lövdén M, et al. LSD-induced entropic brain activity predicts subsequent personality change. Human Brain Mapping. 2016;37:3203–3213. doi:10.1002/hbm.23234.

Margulies DS, Ghosh SS, Goulas A, et al. Situating the default-mode network along a principal gradient of macroscale cortical organization. Proceedings of the National Academy of Sciences. 2016;113:12574– 12579. doi:10.1073/pnas.1608282113.

McCulloch DEW, Olsen AS, Ozenne B, et al. Multi-metric evaluations of acute psychedelic effects on fMRI brain entropy. Nature Communications. 2026;17:7940. doi:10.1038/s41467-026-74215-5.

Novelli L, Stoliker D, Barta T, et al. PsiConnect: Multimodal Neuroimaging of Context-Dependent Brain and Behaviour Dynamics under Psilocybin. Scientific Data. 2026;13:1407. doi:10.1038/s41597-026-07312-1.

Padawer-Curry JA, Krentzman OJ, Kuo CC, et al. Psychedelic 5-HT2A receptor agonism alters neurovascular coupling and differentially affects neuronal and hemodynamic measures of brain function. Nature Neuroscience. 2025;28:2330–2343. doi:10.1038/s41593-025-02069-z.

Prichard D, Theiler J. Generating surrogate data for time series with several simultaneously measured variables. Physical Review Letters. 1994;73:951–954. doi:10.1103/PhysRevLett.73.951.

Raut RV, Snyder AZ, Raichle ME. Hierarchical dynamics as a macroscopic organizing principle of the human brain. Proceedings of the National Academy of Sciences. 2020;117:20890–20897. doi:10.1073/pnas.2003383117.

Richman JS, Moorman JR. Physiological time-series analysis using approximate entropy and sample entropy. American Journal of Physiology Heart and Circulatory Physiology. 2000;278:H2039–H2049. doi:10.1152/ajpheart.2000.278.6.H2039.

Schaefer A, Kong R, Gordon EM, et al. Local-global parcellation of the human cerebral cortex from intrinsic functional connectivity MRI. Cerebral Cortex. 2018;28:3095–3114. doi:10.1093/cercor/bhx179.

Schartner MM, Carhart-Harris RL, Barrett AB, Seth AK, Muthukumaraswamy SD. Increased spontaneous MEG signal diversity for psychoactive doses of ketamine, LSD and psilocybin. Scientific Reports. 2017;7:46421. doi:10.1038/srep46421.

Shinn M, Hu A, Turner L, et al. Functional brain networks reflect spatial and temporal autocorrelation. Nature Neuroscience. 2023;26:867–878. doi:10.1038/s41593-023-01299-3.

Siegel JS, Subramanian S, Perry D, et al. Psilocybin desynchronizes the human brain. Nature. 2024;632:131–138. doi:10.1038/s41586-024-07624-5.

Stoliker D, Novelli L, Khajehnejad M, et al. Psychedelics align brain activity with context. Nature. 2026;656:936–947. doi:10.1038/s41586-026-10910-z.

Subramanian S, Reneau TR, Perry D, et al. Psilocybin’s acute and persistent brain effects: a precision imaging drug trial. Scientific Data. 2025;12:941. doi:10.1038/s41597-025-05189-0.

Tagliazucchi E, Carhart-Harris R, Leech R, Nutt D, Chialvo DR. Enhanced repertoire of brain dynamical states during the psychedelic experience. Human Brain Mapping. 2014;35:5442–5456. doi:10.1002/hbm.22562.

Wang Z, Li Y, Childress AR, Detre JA. Brain entropy mapping using fMRI. PLoS ONE. 2014;9:e89948. doi:10.1371/journal.pone.0089948.

Xiong W, Faes L, Ivanov PCh. Entropy measures, entropy estimators, and their performance in quantifying complex dynamics: Effects of artifacts, nonstationarity, and long-range correlations. Physical Review E. 2017;95:062114. doi:10.1103/PhysRevE.95.062114.

Zang Y, Jiang T, Lu Y, He Y, Tian L. Regional homogeneity approach to fMRI data analysis. NeuroImage. 2004;22:394–400. doi:10.1016/j.neuroimage.2003.12.030.

