## Supplementary for "Psilocybin Reconfigures Cortical Dynamics Across Temporal, Spatial, and Hierarchical Scales"

### Supplementary Methods

#### Dataset and participants

Data were drawn from the publicly available PsiConnect dataset (OpenNeuro ds006110), an open-label multimodal study of psychedelic-naive adults scanned at baseline and after oral psilocybin administration (19 mg). Acute imaging began approximately 80 min after dosing; the eight-minute eyes-closed resting-state run was acquired first in a fixed task order on a 3 T Siemens Skyra and contained 505 volumes (TR = 0.91 s). Fifty-seven participants had paired baseline (ses-01) and post-psilocybin (ses-02) resting-state scans that passed the specified motion criterion: no more than 20% of volumes with framewise displacement greater than 0.5 mm in either session. This secondary analysis is a within-participant pre/post comparison without a placebo condition.

#### Image preprocessing and denoising

We used fMRIPrep 22.0.2 (Esteban et al., 2019) preprocessed BOLD images in MNI152NLin2009cAsym space and the corresponding confound files. Confound regression included the Friston-24 expansion of six rigid-body motion estimates and the original, derivative, squared, and squared-derivative terms for global, white-matter, and cerebrospinal-fluid signals, together with an intercept and linear trend. Volumes with framewise displacement greater than 0.5 mm were interpolated after regression. Unless otherwise stated, signals were band-pass filtered at 0.01-0.08 Hz and standardized voxelwise. The first four cleaned volumes were discarded, leaving at most 501 analyzed time points per run. Analyses were restricted to a common gray-matter mask defined by tissue probability at least 0.30 and coverage in at least 80% of participants. Spatial analyses used 8-mm FWHM smoothing within the fMRIPrep brain mask; ReHo was calculated before smoothing and its final map was then smoothed by 8 mm. Although PsiConnect used multi-echo acquisition, this analysis used the fMRIPrep preprocessed BOLD series rather than a tedana-denoised multi-echo derivative.

#### Dynamic measures

BEN was quantified as sample entropy with embedding dimension m = 3 and tolerance r = 0.6 times each voxel's temporal standard deviation, excluding self-matches (Richman & Moorman, 2000, Wang et al., 2014). Lempel-Ziv complexity used median binarization and the normalized LZ76 estimator. DFA exponents were fitted over windows of 4, 8, 16, 32, and 64 samples and interpreted as scaling summaries over this finite range. MSSD was the mean squared difference between successive standardized samples. ALFF was the mean Fourier amplitude from 0.01 to 0.08 Hz, and fALFF was its fraction of total nonzero-frequency amplitude (Zou et al., 2008). These amplitude measures used nuisance residuals without band-pass filtering or voxelwise variance normalization, and were normalized by the gray-matter mean (mALFF and mfALFF). ReHo was Kendall's W across each voxel and its 26 neighbors (Zang et al., 2004). Intrinsic timescale was the unbiased BOLD autocorrelation integral to its first zero crossing; this operational measure is distinct from electrophysiological neuronal time constants (Raut et al., 2020).

#### Global spatial measures

Spatial BEN was calculated within each BOLD frame from contiguous four-voxel templates along each of the three voxel axes. Match counts for three- and four-voxel prefixes used a maximum absolute difference threshold of 0.6 times the within-frame spatial standard deviation. Self-matches were excluded. We sampled 131,072 template pairs per axis, reused the same draws across conditions within each participant, and averaged the negative log ratio of four- to three-voxel matches over frames and axes. NGSC was the normalized Shannon entropy of the eigenvalue spectrum of the voxelwise correlation matrix (Siegel et al., 2024). Both measures used nuisance-cleaned, 8-mm-smoothed gray-matter BOLD. In ds006072 we restricted both measures to each participant’s common valid voxel support across the three conditions. These global summaries were not used as predictor features.

#### Functional hierarchy and traveling waves

Resting-state time series were summarized using the Schaefer-100 seven-network atlas (Schaefer et al., 2018). Pearson functional-connectivity matrices were Fisher-z transformed. Diffusion-map embedding used the top 10% of each row, normalized-angle affinity, alpha = 0.5, and three components, followed by Procrustes alignment to a reference derived from pooled baseline and post-dose connectivity matrices. The component showing the strongest sensory-to-transmodal separation (Margulies et al., 2016) was treated as the hierarchy gradient. Motivated by the organization of large-scale phase propagation (Raut et al., 2021), exploratory hierarchy-aligned waves were estimated after ordering parcels into ten bins along this gradient. Candidate waves required phase-gradient regression R2 at least 0.5, adjacent-step coherence at least 0.7, and absolute circular phase span at least pi/2. Because phase slope was defined across ordered bins rather than physical distance, it indexes phase organization and not propagation velocity.

#### Covariance, similarity, and structure-function analyses

At the Desikan-68 scale, group covariance networks were defined as across-participant Spearman correlations for each functional metric separately at baseline and after psilocybin. In adjusted analyses, the subjectwise cortex-wide metric mean was removed by parcelwise across-participant linear regression before covariance estimation. Session-label permutation (10,000 iterations) tested edge and network-pattern changes. Functional-similarity networks were constructed within each participant/session as cosine similarity between parcels after feature-wise z-scoring of eight functional measures across parcels. Morphometric-similarity networks used cortical thickness, surface area, and gray-matter volume. Sparse topology retained the strongest 20% of positive edges. Structure-function coupling was the within-participant Spearman correlation across 68 parcels between each morphometric feature and each functional metric; these analyses were treated as exploratory because they do not constitute white-matter structural connectivity.

#### Molecular and behavioral associations

Voxelwise post-minus-pre mean effect maps were parcellated with Schaefer-400 and compared using Spearman correlations with 21 PET-derived molecular maps and seven Allen Human Brain Atlas cell-type marker maps (Hansen et al., 2022, Markello et al., 2022). The panel includes receptors, transporters, SV2A, and TSPO. ReHo used the mean-FD-change-adjusted effect map. Valid effects covered 272 or 273 parcels. The original molecular analysis used 10,000 hemisphere-preserving centroid rotations of effect maps, with pairwise valid parcels and two-sided deviations relative to the null mean. BH-FDR was evaluated within effect families and across all 224 comparisons. A sensitivity analysis retained each observed effect mask, rotated complete molecular maps, recomputed ranks within each valid subset, and used absolute two-sided null correlations with plus-one p-value correction; FDR again covered 224 comparisons. Centroid rotations provide an approximate spatial null. Exploratory brain-behavior correlations used uncorrected p < 0.05.

#### Statistical inference

Paired t-tests quantified pre/post changes. Voxelwise p values were controlled with Benjamini-Hochberg FDR within each gray-matter mask and metric. Parcel tests were corrected across the number of parcels, edge tests within the 2,278 Desikan edges, and network-pair tests across 28 pairs. Unless explicitly labeled exploratory, q < 0.05 was considered significant.

#### Independent cohort and matched duration analysis

We analyzed one baseline, one acute psilocybin and one acute methylphenidate resting run for each of seven participants from the independent randomized crossover dataset ds006072 (Siegel et al., 2024, Subramanian et al., 2025). We selected BOLDREST1 run 1 and used the first baseline except for P5, whose ses-3 baseline matched the phase-encoding direction of both acute scans. Open-label repeat visits were not included. Multi-echo BOLD was processed with fMRIPrep 25.2.5, including opposite phase-encoding distortion correction, and resampled onto the PsiConnect analysis grid. We used the same nuisance, smoothing and metric definitions with the native repetition time of 1.761 s. Within each participant and metric, condition means were computed over the intersection of valid gray-matter voxels across the three runs.

Full-length analyses used all available samples after the initial four-volume discard. The duration sensitivity analysis recomputed each metric using the first 204 retained frames (approximately six minutes), matching the shortest run. Nuisance regression and filtering were performed before window selection using the original metric-specific processing order; this analysis matches metric input length rather than fitting preprocessing anew on the short window. BEN used m = 3 and r = 0.6 times the temporal standard deviation of the analyzed series. We report paired difference signs for both drug-to-baseline contrasts and the direct psilocybin-minus-methylphenidate contrast.

We averaged valid positive metric values within each Schaefer-400 parcel and applied frozen logistic regression pipelines fitted to the 57 PsiConnect pairs. Imputation, scaling and coefficients were fixed before external scoring. We report within-person condition orderings, not thresholded classifications or calibrated probabilities. Counts of directional and ordering agreement are descriptive; the participant, not the run, is the unit of comparison.

#### Cross cohort cortical topography

We compared mean PsiConnect post-minus-baseline maps with external mean paired contrasts on the Schaefer-400 atlas. External contrasts were psilocybin minus baseline, methylphenidate minus baseline, and psilocybin minus methylphenidate, each at full length and 204 frames. We computed Spearman correlations within jointly covered parcels. Ten thousand hemisphere-preserving spatial permutations rotated the external values and their coverage together; correlations were recomputed on each valid overlap. Two-sided spin p values were BH-adjusted separately for each contrast across both lengths and all measures within each family (metric maps or FC gradients). Discovery spatial maps were compared with external parcel means; the external participants, not parcels, supplied the paired effects.

For FC gradients, we extracted atlas-matched time series for all 21 external scans and retained 396 parcels common to both cohorts. We ran the same three-component diffusion embedding on Fisher-z Pearson connectivity from both cohorts, keeping the strongest 10% of each row and using normalized-angle affinity. Individual embeddings were aligned by orthogonal Procrustes to the fixed PsiConnect reference, formed from both discovery sessions. The external time series were analyzed at full length and over the first 204 retained frames. We compared maps of changes in aligned gradient coordinates, rather than correlations between metric changes and a static cortical hierarchy.

Separately, we tested PsiConnect effect maps against an independent sensory-to-transmodal hierarchy using spatial permutations and FDR within the eight-effect family. Partial rank correlations with CB1 and mGluR5 adjusted for this hierarchy, with spatial nulls and FDR across 16 effect-by-receptor tests. PET templates represent population-average reference maps, not receptor measurements in either cohort.

### Supplementary Results

#### Regional BOLD entropy increased after psilocybin

Voxelwise analysis identified 1,397 of 22,740 gray-matter voxels with FDR-significant paired BEN changes, comprising 1,396 increases and one decrease (main-text Fig. 1a). Significant increases occurred across default-mode (489 voxels), visual (297), control (223), dorsal-attention (200), and salience/ventral-attention (132) territories. These counts describe the distribution of detected effects and are not normalized by network size. The result establishes a regional increase in BOLD temporal irregularity under the specified sample-entropy parameters. Of these, 1,340 positive voxels fell within the Schaefer-400 surface display in main-text Fig. 1a. The voxel counts in Fig. 1b include the remaining 57 significant voxels, including the single decrease.

#### Local synchrony and temporal persistence decreased

The greatest extent of significant voxels was observed for a reduction in ReHo: 18,338 of 22,622 voxels survived FDR, with 18,083 decreases. This remained extensive after adjustment for mean framewise-displacement change (18,495 significant voxels; 18,207 decreases). MSSD increased in 9,649 voxels, indicating enhanced moment-to-moment signal change. DFA decreased in 7,678 voxels, consistent with altered scaling over the fitted window range. Intrinsic timescale shortened by an average of 0.136 s (paired p = 1.41 x 10^-5), with FDR-significant decreases in 4,320 of 22,622 voxels. LZC increased in 250 voxels. mALFF and mfALFF showed bidirectional spatial changes, indicating redistribution rather than a unitary amplitude effect.

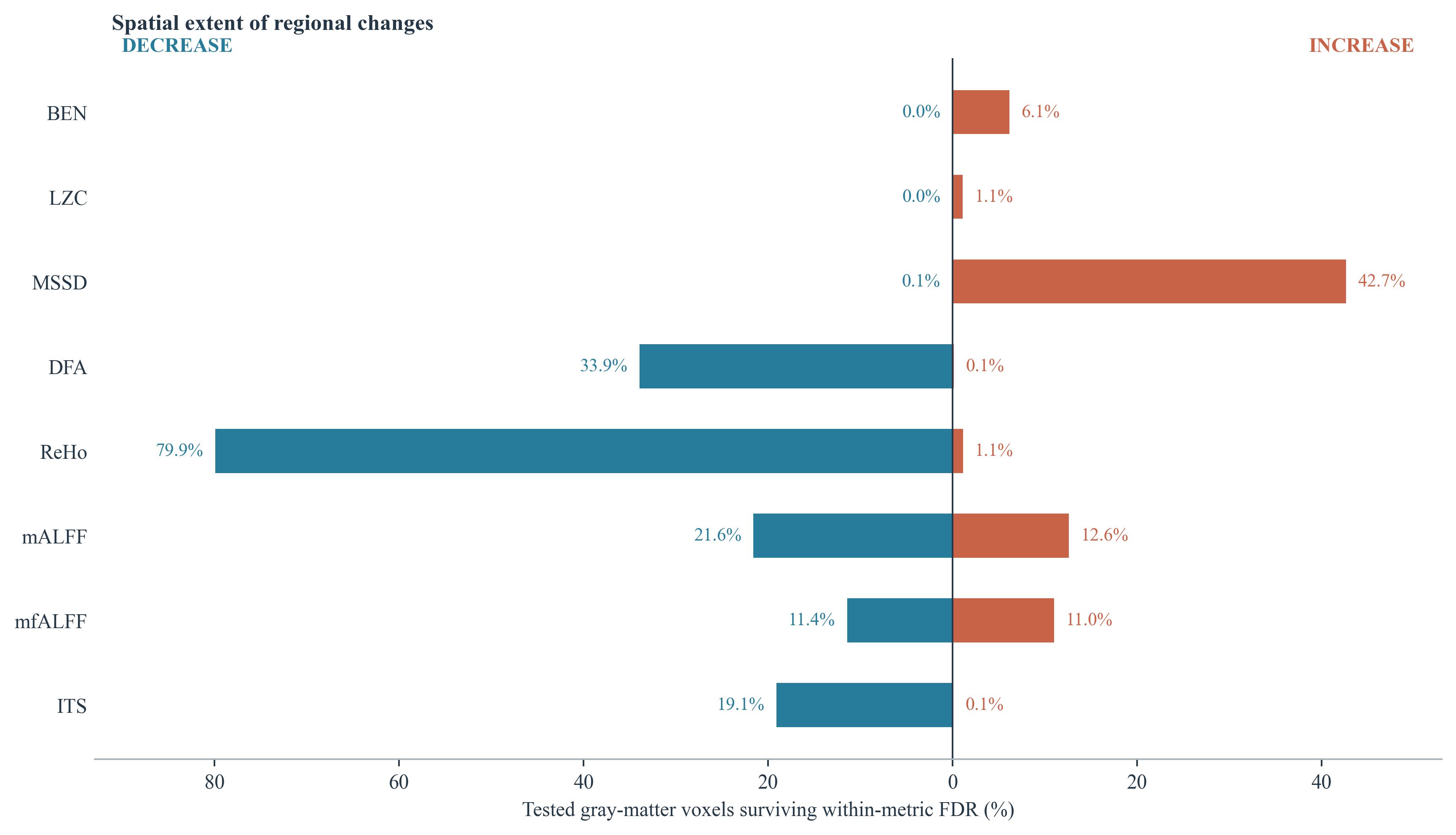

**Supplementary Figure 1.** Direction and spatial extent of FDR-significant voxelwise effects. Bar length is the percentage of tested gray-matter voxels with significant increases or decreases. Metrics differ in scale and should not be compared by raw effect magnitude.

#### Spatial pattern irregularity in PsiConnect

Global spatial BEN increased by 0.0242 after psilocybin (95% CI 0.0177 to 0.0306; paired p = 4.86 × 10⁻¹⁰), with increases in 49 of 57 participants. Voxelwise NGSC increased by 0.0192 (95% CI 0.0148 to 0.0235; paired p = 3.16 × 10⁻¹²; 49 of 57 participants). Thus, within-frame spatial templates were less likely to remain similar when extended, and variance was less concentrated across spatial modes.

#### Surrogate comparisons and preprocessing sensitivity

Main-text Fig. 2 shows the full-sample model estimates, the distance-model contrast by sample and smoothing, and preprocessing sensitivity. For the spatial comparison, common phase minus independent phase was 0.02498 (95% CI 0.01832 to 0.03164) in the full sample. Common phase minus zero-lag covariance was 0.00010 (95% CI −0.00002 to 0.00021). For temporal BEN, independent phase minus covariance was 0.00532 (95% CI 0.00371 to 0.00693). All estimates use participant-level paired post-minus-baseline contrasts, averaging 10 realizations within each scan before inference. The no-GSR comparisons use the same 49 participants in both pipelines.

#### External changes and sensitivity to scan duration

ReHo was lower after psilocybin than at baseline in all seven external participants with both full-length and 204-frame analyses (mean paired changes −0.0299 and −0.0269, respectively). The frozen ReHo model also assigned higher scores to psilocybin than baseline in all seven participants at both durations. Across all three conditions, six of seven participants retained the identical ReHo-model ordering after duration matching. These results show agreement across cohorts in the baseline contrast, both for the gray-matter mean and for the parcelwise signature.

Spatial BEN increased after psilocybin relative to baseline in all seven external participants at both durations (mean changes +0.0271 at full length and +0.0235 at 204 frames). Methylphenidate also increased it in six of seven at both durations (+0.0106 and +0.0086). Psilocybin exceeded methylphenidate in five of seven at full length and four of seven at 204 frames; the corresponding mean differences were +0.0165 and +0.0150.

NGSC increased after psilocybin relative to baseline in six of seven at each duration (+0.0128 and +0.0148), although only five retained the same individual direction. After methylphenidate, NGSC increased in five of seven at full length and four of seven at 204 frames (+0.0046 and +0.0019). Psilocybin exceeded methylphenidate in four of seven and five of seven, respectively (+0.0082 and +0.0130). These comparisons were descriptive and neither spatial measure entered the frozen models.

The active-comparator contrast was less uniform. Mean ReHo was lower after psilocybin than methylphenidate in five of seven participants at full length and four of seven at 204 frames; six individuals retained their contrast direction. The mean psilocybin-minus-methylphenidate difference was −0.0170 and −0.0135, respectively. Methylphenidate itself reduced ReHo relative to baseline in the same five participants at both durations. In contrast to the mean ReHo comparison, the frozen ReHo model ranked psilocybin above methylphenidate in five participants at full length and six at 204 frames. Because the model weights regional ReHo patterns, its ordering need not match the direction of a whole-gray-matter mean. These descriptive results do not establish drug specificity.

On metric-specific valid voxel support, BEN increased after psilocybin in the same six participants at both durations, with mean changes of −0.0002 and +0.0088. BEN also increased after methylphenidate in seven participants at full length and six at 204 frames. Intrinsic timescales shortened after psilocybin relative to baseline in five participants at full length and seven after duration matching; the direct drug comparison retained its direction in four of seven. Supplementary Table 1 reports the direction and ordering checks for all measures. BEN-only and three-metric model scores were nearly saturated, so their raw score scale should not be interpreted as calibrated evidence. This full-length and duration-matched analysis used its metric-specific valid-voxel rule. The fixed common-support surrogate analysis in main-text Fig. 3 applied a shared mask across all three visits and signal types. On that common support at 204 frames, observed BEN increased in seven of seven participants (mean +0.00793).

#### External cohort duration sensitivity

**Supplementary Table 1.** Agreement between full-length and 204-frame analyses in seven participants.

| **Measure or frozen model** | **PSIL vs baseline** | **MTP vs baseline** | **PSIL vs MTP** |
| --- | --- | --- | --- |
| Mean ReHo | 7 | 7 | 6 |
| Mean BEN | 7 | 6 | 6 |
| Mean intrinsic timescale | 5 | 6 | 4 |
| ReHo model | 7 | 7 | 6 |
| BEN model | 7 | 7 | 5 |
| Intrinsic timescale model | 6 | 4 | 5 |
| Three-metric model | 6 | 7 | 5 |
| ReHo and timescale model | 6 | 6 | 7 |
| Global spatial BEN | 7 | 7 | 6 |
| Global NGSC | 5 | 6 | 6 |

Entries are counts out of seven with the same sign (mean metrics) or within-participant pairwise score ordering (frozen models) in both analyses. PSIL, psilocybin; MTP, methylphenidate.

#### Regional reproducibility of temporal changes

In PsiConnect, the largest overlaps of FDR-detected BEN increases included default-network prefrontal and posterior cingulate parcels and visual cortex. Timescale decreases had prominent overlap in bilateral posterior cingulate and lateral and medial prefrontal control territories; DFA decreases also involved default and control areas. These are atlas localizations of voxelwise effects rather than parcel-level significance tests or network enrichment estimates.

Main-text Fig. 4 shows the BEN, intrinsic timescale and aligned gradient maps. Beyond those measures, MSSD, DFA, LZC, mALFF and mfALFF showed positive cross-cohort correspondence at both durations for psilocybin minus baseline. The external psilocybin-minus-methylphenidate maps also showed positive correspondence with the PsiConnect maps at both durations for timescale, DFA, MSSD, mALFF and mfALFF. BEN passed correction for that contrast at full length. Spatial correspondence quantifies the similarity of regional effect patterns.

#### Connectivity differentiation contracted after psilocybin

The second connectivity gradient was the component most strongly aligned with the sensory-to-transmodal hierarchy. Its range decreased after psilocybin (p = 7.94 x 10^-4, q = 0.0024), as did its variance (p = 1.41 x 10^-4, q = 8.45 x 10^-4); 24 of 100 parcels showed FDR-significant loading changes. Among seven traveling-wave metrics, median absolute phase slope decreased (p = 0.0066, q = 0.046), whereas wave prevalence did not change. These results suggest compression of hierarchical differentiation together with a shallower phase gradient across ordered hierarchy bins, rather than disappearance of traveling-wave events. They do not by themselves establish slower physical propagation.

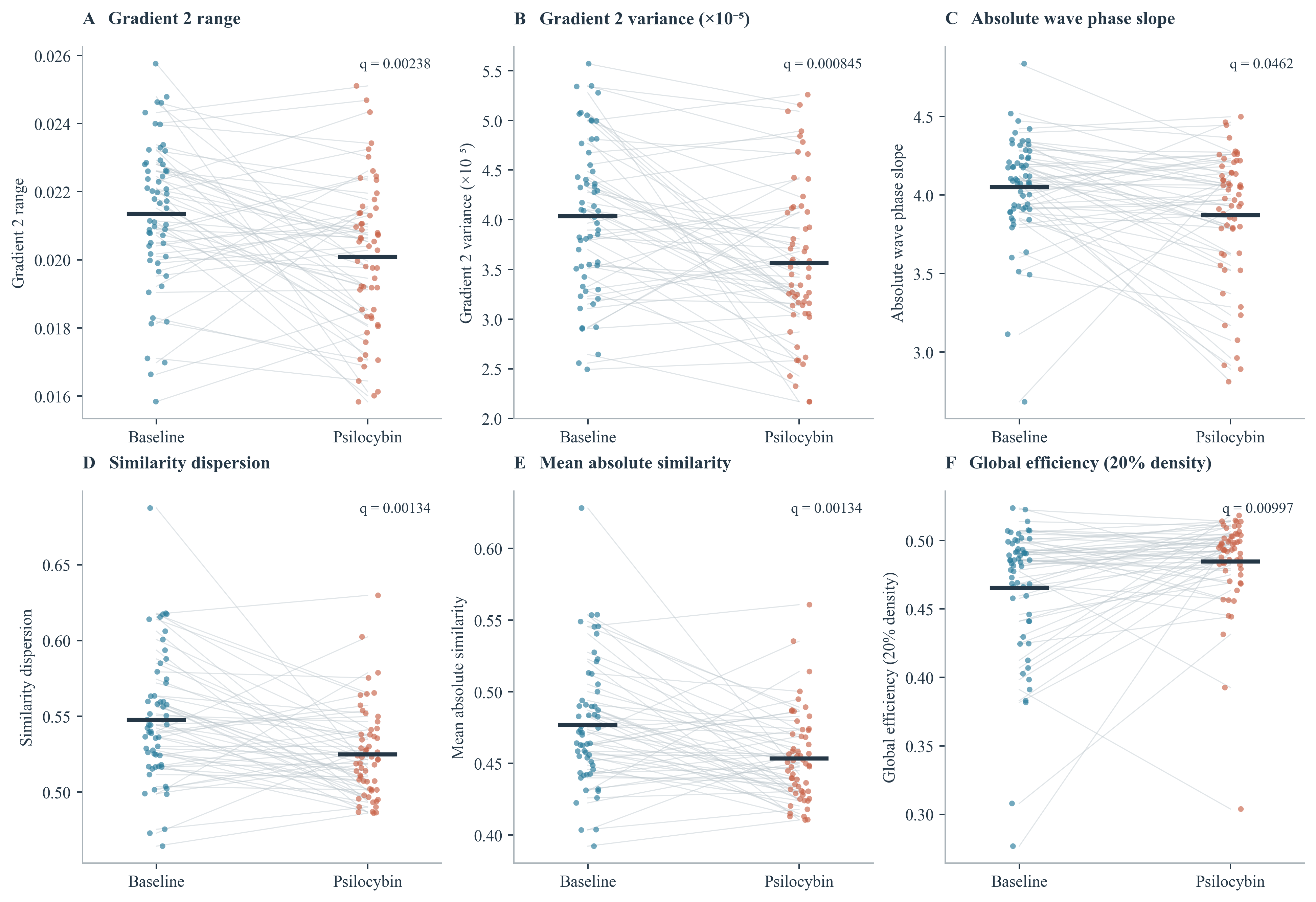

**Supplementary Figure 2.** Paired changes in hierarchical organization and functional-similarity topology. (A,B) Range and variance of aligned connectivity gradient 2. (C) Absolute hierarchy-aligned wave phase slope. (D-F) Functional-similarity dispersion, mean absolute similarity, and global efficiency at the selected 20% positive-edge density. Points show individual participants; gray lines connect paired sessions; dark horizontal segments mark means. q values refer to the original six-gradient, seven-wave, or five-topology test families. Phase slope does not measure physical propagation speed.

#### Cross cohort changes in FC gradient coordinates

Among aligned gradient changes for the external psilocybin-minus-baseline contrast, G3 showed positive discovery correspondence at both durations (rho = 0.228 and 0.263; spin q = 0.0423 and 0.0090). G1 and G2 passed correction at 204 frames (rho = 0.160 and 0.323; q = 0.0274 and 0.00060), but not at full length (q = 0.165 and 0.237). Methylphenidate-minus-baseline also showed G3 correspondence at both durations (rho = 0.477 and 0.475; both q = 0.00030), numerically larger than the psilocybin baseline correspondence. No component passed correction for the direct drug contrast at full length; at 204 frames, G1 correspondence was positive (rho = 0.183, q = 0.0216) and G3 correspondence was negative (rho = -0.273, q = 0.0357). Component labels refer to the common PsiConnect reference, so G3 should not be equated with the sensory-to-transmodal component G2.

#### Regional functional profiles showed altered network organization

Functional-similarity networks showed lower similarity dispersion (Cohen's dz = -0.50, q = 0.0013) and lower mean absolute similarity (dz = -0.49, q = 0.0013), accompanied by higher global efficiency at the specified strongest-20%-positive-edge density (dz = 0.38, q = 0.010). Fifty-two of 2,278 edges changed after FDR. Mean coupling between morphometric- and functional-similarity networks did not change (mean rho 0.118 to 0.113, p = 0.585), providing no detected difference in this sample; preservation was not tested using an equivalence design. Network-efficiency inference remains conditional on the chosen edge density.

#### Global adjustment revealed lower amplitude and synchrony covariance coupling

After parcelwise residualization of the subjectwise cortex-wide metric mean, the correspondence between mALFF and ReHo covariance-edge patterns decreased from rho = 0.793 to 0.551 (10,000-permutation p = 0.00030, FDR q = 0.0084 across 28 network pairs; Supplementary Figure 3B). The association concerns regional covariance after global adjustment. Without that adjustment, mALFF-ReHo coupling changed from 0.576 to 0.515 (p = 0.272), and no network pair survived FDR. Thus, the detected coupling change depends on separating regional covariance from cortex-wide metric variation.

#### CB1 and mGluR5 distributions aligned with temporal effects

Six spatial associations survived FDR across all 224 molecular tests. CB1 density was positively associated with MSSD and LZC effects and negatively associated with DFA and intrinsic-timescale effects. mGluR5 density was positively associated with MSSD and negatively associated with DFA. BEN showed a negative association with the 5-HT1A map that survived within-BEN-family correction (rho = -0.419, pspin = 0.0017, qwithin = 0.0357) but narrowly missed correction across all tests (qall = 0.053). No cell-type association survived across-test FDR. All six associations remained significant in the fixed-mask sensitivity analysis. That analysis additionally identified LZC-mGluR5 (rho = 0.335, q = 0.0261) and timescale-mGluR5 (rho = -0.315, q = 0.0392) associations. Supplementary Figure 3A marks the original six associations, preserving the distinction between the original test and its sensitivity analysis. No PsiConnect metric effect passed FDR against the independent cortical hierarchy; for BEN, rho = 0.026 and spin p = 0.839. After controlling for hierarchy across approximately 272 covered parcels, higher CB1 and mGluR5 reference density remained associated with greater MSSD and LZC increases and stronger DFA and timescale decreases (q < 0.05 across the 16 partial tests). BEN did not survive this correction for either receptor (q = 0.083 and 0.065). Supplementary Figure 4 shows the hierarchy and hierarchy-adjusted receptor associations.

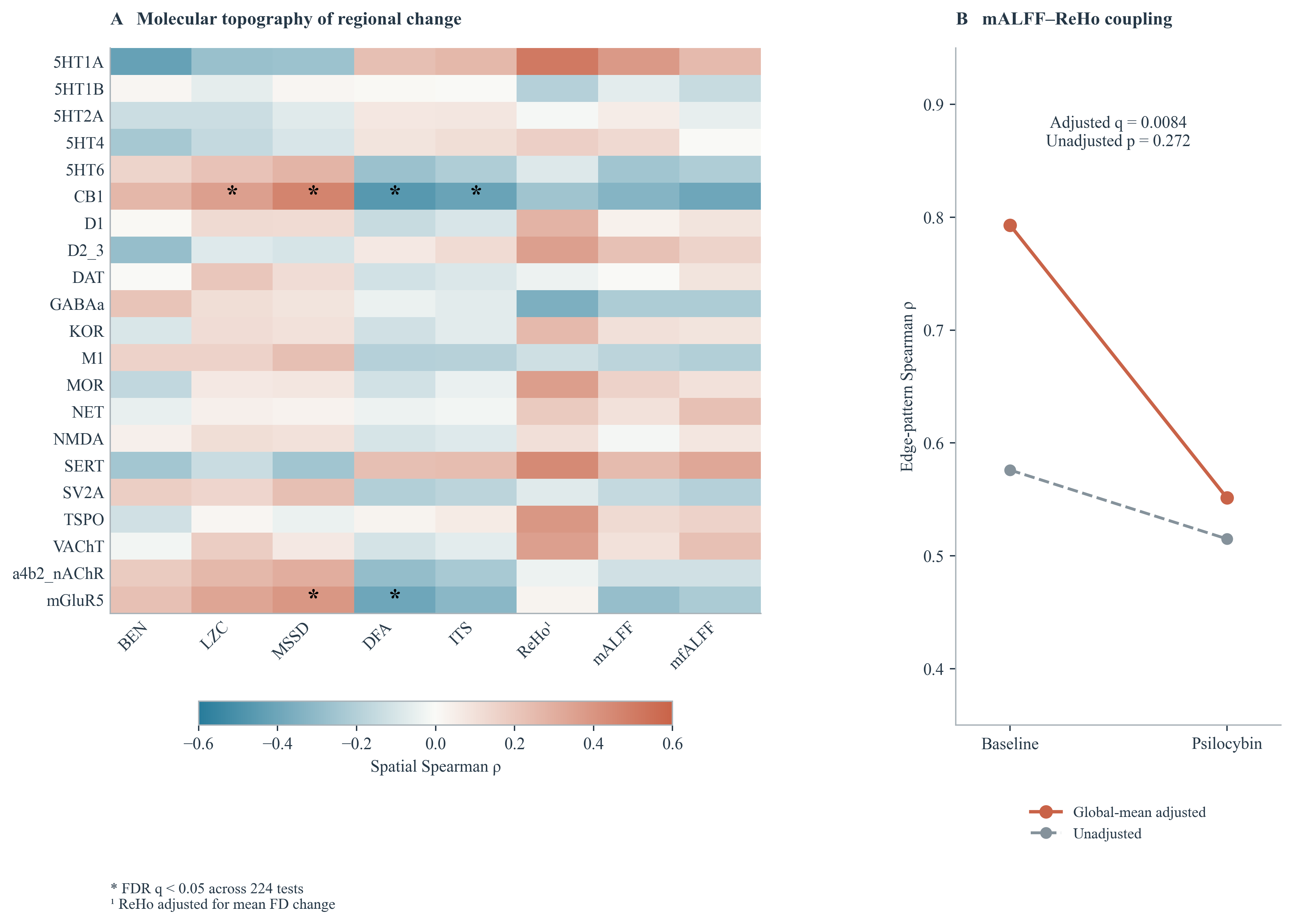

**Supplementary Figure 3.** Molecular topography and cross-metric covariance coupling. (A) Spatial Spearman correlations between eight regional effect maps and all 21 PET molecular maps. Asterisks mark the six associations surviving FDR across the full 224-test molecular and cell-type family. ReHo effects are adjusted for mean FD change. (B) mALFF-ReHo covariance-edge coupling with and without subjectwise global-mean residualization. Points represent group network estimates, not individual participants; significance is based on session-label permutation. All six marked associations also survived the fixed-mask sensitivity analysis.

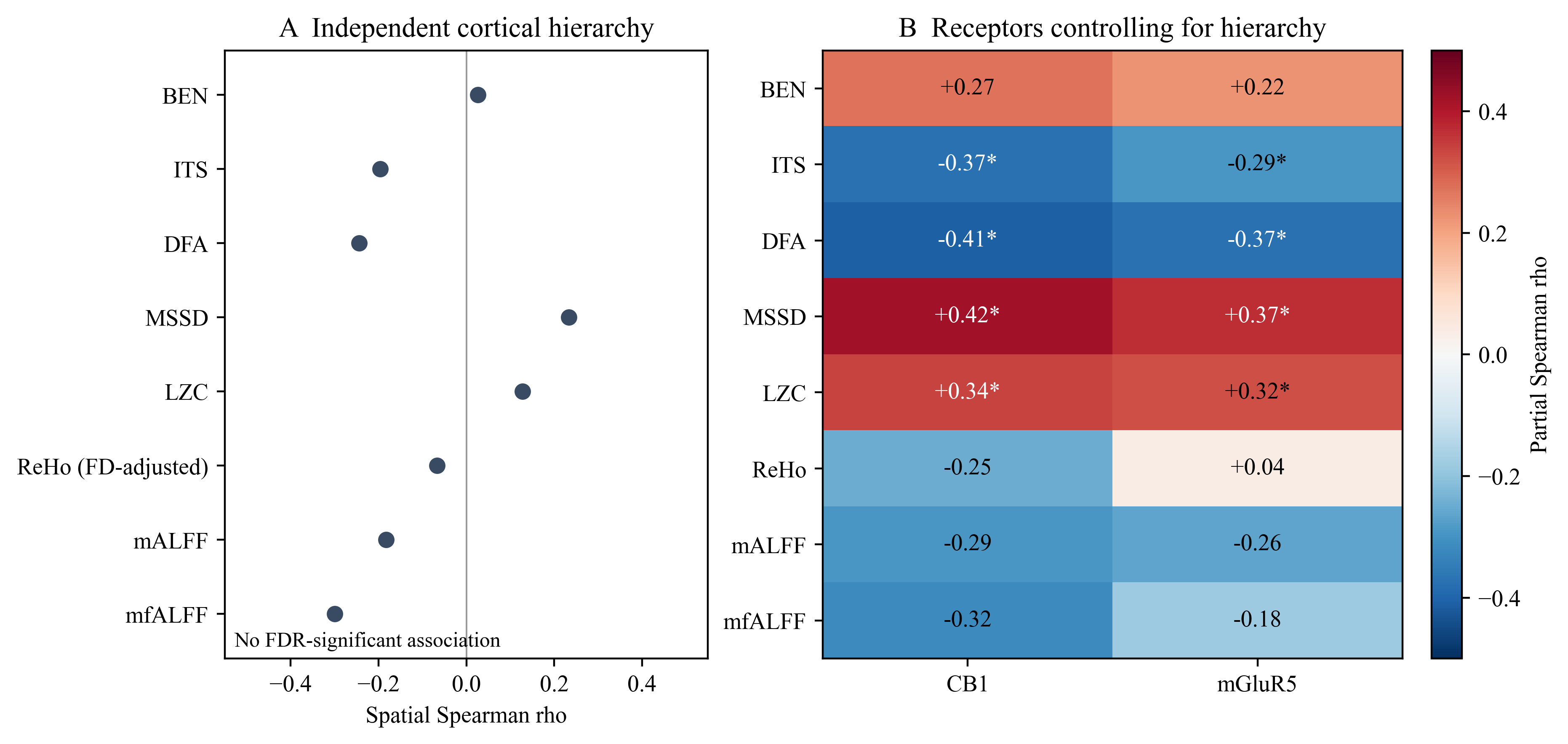

**Supplementary Figure 4.** Spatial associations of PsiConnect effect maps with an independent cortical hierarchy and receptor reference maps. (A) No hierarchy correlation survived FDR across eight effect maps. (B) Partial Spearman correlations with CB1 and mGluR5 controlling for the independent hierarchy. Asterisks indicate spatial-null q < 0.05 across 16 tests. ReHo was adjusted for framewise displacement in both panels. Receptor associations reflect spatial colocalization and do not demonstrate mediation.

#### Exploratory associations between individual changes

We used Spearman correlations to relate post-minus-pre changes in gray-matter mean intrinsic timescale and regional homogeneity to each other and to the range and variance of connectivity gradient 2 (57 paired participants, five tests, BH-FDR). The largest correlations were between changes in regional homogeneity and gradient variance (rho = 0.265, p = 0.047, q = 0.135) and between changes in intrinsic timescale and regional homogeneity (rho = 0.257, p = 0.054, q = 0.135). No association survived correction.

#### Whole gray matter BEN summary

A separately generated whole-gray-matter BEN summary showed no detected session difference (mean change = -0.00027, paired t = -0.19, p = 0.847). This summary and the smoothed common-mask voxelwise analysis arose from separate processing branches and do not constitute a matched-pipeline comparison of global and regional effects.

#### Exploratory morphometric-functional coupling

Supplementary Table 2 summarizes the group-level morphometric and functional covariance comparisons.

**Supplementary Table 2.** Strongest exploratory changes in parcelwise morphometric-functional coupling. In keeping with the exploratory analysis plan, p values are not additionally corrected across the 24 metric combinations; ROI-wise analyses elsewhere are corrected across the number of parcels. These are spatial-profile correlations, not tractography-based structural connectivity.

| **Morphometric feature** | **Functional metric** | **rho pre** | **rho post** | **Delta Fisher z** | **Uncorrected p** |
| --- | --- | --- | --- | --- | --- |
| thickness | ReHo | -0.416 | -0.300 | 0.134 | 1.54e-05 |
| surface_area | mALFF | -0.014 | -0.124 | -0.111 | 9.3e-05 |
| thickness | BEN | 0.230 | 0.061 | -0.173 | 0.000153 |
| gray_volume | mALFF | -0.075 | -0.157 | -0.083 | 0.000499 |
| thickness | mALFF | -0.261 | -0.114 | 0.153 | 0.00122 |
| surface_area | ReHo | 0.447 | 0.399 | -0.058 | 0.00724 |
| thickness | MSSD | 0.208 | 0.098 | -0.112 | 0.0113 |
| thickness | ITS | -0.111 | -0.006 | 0.106 | 0.0178 |
| thickness | DFA | -0.191 | -0.100 | 0.093 | 0.024 |
| thickness | LZC | 0.162 | 0.079 | -0.084 | 0.0499 |

#### Nominal behavioral associations

Whole-gray-matter BEN change showed nominal negative associations with blissful unity (rho = -0.321, p = 0.0216), blissfulness (rho = -0.284, p = 0.0394), and a composite associative/blissful-unity/spiritual score (rho = -0.289, p = 0.0395). Other nominal findings linked gradient and wave changes to ineffability, unity, positive mood, and oceanic boundlessness. These correlations were not corrected for the full imaging-by-scale search and should not be interpreted as confirmed brain-behavior relationships. Voxelwise BEN correlations and time-by-scale interactions yielded no FDR-significant voxels.

**Supplementary Table 3.** First 15 nominal imaging-behavior associations sorted by uncorrected p. No multiplicity correction was applied; these findings are exploratory.

| **Imaging change** | **Scale** | **n** | **Spearman rho** | **Uncorrected p** |
| --- | --- | --- | --- | --- |
| G2_variance_change | MEQ30_INEFFABILITY | 57 | -0.355 | 0.0067 |
| median_abs_phase_slope_change | MEQ30_INEFFABILITY | 57 | -0.348 | 0.0080 |
| median_abs_phase_slope_change | ASC11_BLISSFUL | 53 | -0.335 | 0.0141 |
| wave_fraction_change | ASC11_COMPOSITE_SENSORY_AUDIOVISUAL_COMPLEX_ELEMENTARY | 50 | 0.343 | 0.0149 |
| median_abs_phase_slope_change | MEQ30_TRANSCEND | 57 | -0.319 | 0.0155 |
| median_abs_phase_slope_change | MEQ30_POSITIVE | 57 | -0.314 | 0.0174 |
| sensory_to_transmodal_fraction_change | ASC11_UNITY | 51 | -0.330 | 0.0179 |
| G2_variance_change | ASC5_OCEANICBOUNDLESS | 50 | -0.333 | 0.0182 |
| sensory_to_transmodal_fraction_change | ASC11_COMPOSITE_BLISSFUL_UNITY | 51 | -0.324 | 0.0205 |
| BEN_change | ASC11_COMPOSITE_BLISSFUL_UNITY | 51 | -0.321 | 0.0216 |
| G2_variance_change | ASC11_COMPOSITE_BLISSFUL_UNITY | 51 | -0.315 | 0.0244 |
| wave_fraction_change | ASC11_BLISSFUL | 53 | -0.307 | 0.0253 |
| G1_variance_change | ASC11_ANXIETY | 52 | -0.309 | 0.0257 |
| G2_variance_change | ASC11_COMPOSITE_ASSOCIATIVE_BLISFFUL_UNITY_SPIRITUAL | 51 | -0.311 | 0.0261 |
| wave_fraction_change | ASC11_COMPOSITE_BLISSFUL_UNITY | 51 | -0.309 | 0.0274 |

### Supplementary References

Esteban O, Markiewicz CJ, Blair RW, et al. fMRIPrep: a robust preprocessing pipeline for functional MRI. Nature Methods. 2019;16:111-116. doi:10.1038/s41592-018-0235-4.

Hansen JY, Shafiei G, Markello RD, et al. Mapping neurotransmitter systems to the structural and functional organization of the human neocortex. Nature Neuroscience. 2022;25:1569-1581. doi:10.1038/s41593-022-01186-3.

Margulies DS, Ghosh SS, Goulas A, et al. Situating the default-mode network along a principal gradient of macroscale cortical organization. Proceedings of the National Academy of Sciences. 2016;113:12574–12579. doi:10.1073/pnas.1608282113.

Markello RD, Hansen JY, Liu ZQ, et al. Neuromaps: structural and functional interpretation of brain maps. Nature Methods. 2022;19:1472–1479. doi:10.1038/s41592-022-01625-w.

Raut RV, Snyder AZ, Mitra A, et al. Global waves synchronize the brain's functional systems with fluctuating arousal. Science Advances. 2021;7:eabf2709. doi:10.1126/sciadv.abf2709.

Raut RV, Snyder AZ, Raichle ME. Hierarchical dynamics as a macroscopic organizing principle of the human brain. Proceedings of the National Academy of Sciences. 2020;117:20890–20897. doi:10.1073/pnas.2003383117.

Richman JS, Moorman JR. Physiological time-series analysis using approximate entropy and sample entropy. American Journal of Physiology Heart and Circulatory Physiology. 2000;278:H2039-H2049. doi:10.1152/ajpheart.2000.278.6.H2039.

Schaefer A, Kong R, Gordon EM, et al. Local-global parcellation of the human cerebral cortex from intrinsic functional connectivity MRI. Cerebral Cortex. 2018;28:3095–3114. doi:10.1093/cercor/bhx179.

Siegel JS, Subramanian S, Perry D, et al. Psilocybin desynchronizes the human brain. Nature. 2024;632:131-138. doi:10.1038/s41586-024-07624-5.

Subramanian S, Reneau TR, Perry D, et al. Psilocybin’s acute and persistent brain effects: a precision imaging drug trial. Scientific Data. 2025;12:941. doi:10.1038/s41597-025-05189-0.

Wang Z, Li Y, Childress AR, Detre JA. Brain entropy mapping using fMRI. PLoS ONE. 2014;9:e89948. doi:10.1371/journal.pone.0089948.

Zang Y, Jiang T, Lu Y, He Y, Tian L. Regional homogeneity approach to fMRI data analysis. NeuroImage. 2004;22:394-400. doi:10.1016/j.neuroimage.2003.12.030.

Zou QH, Zhu CZ, Yang Y, et al. An improved approach to detection of amplitude of low-frequency fluctuation (ALFF) for resting-state fMRI: fractional ALFF. Journal of Neuroscience Methods. 2008;172:137-141. doi:10.1016/j.jneumeth.2008.04.012.
